# Heterochromatin protein 1 (HP1) is intrinsically required for post-transcriptional regulation of *Drosophila* Germline Stem Cell (GSC) maintenance

**DOI:** 10.1101/474833

**Authors:** Assunta Maria Casale, Ugo Cappucci, Laura Fanti, Lucia Piacentini

## Abstract

A very important open question in stem cells regulation is how the fine balance between GSCs self-renewal and differentiation is orchestrated at the molecular level. In the past several years much progress has been made in understanding the molecular mechanisms underlying intrinsic and extrinsic controls of GSC regulation but the complex gene regulatory networks that regulate stem cell behavior are only partially understood. HP1 is a dynamic epigenetic determinant mainly involved in heterochromatin formation, epigenetic gene silencing and telomere maintenance. Furthermore, recent studies have revealed the importance of HP1 in DNA repair, sister chromatid cohesion and, surprisingly, in positive regulation of gene expression. Here, we show that HP1 plays a crucial role in the control of GSC homeostasis in *Drosophila*. Our findings demonstrate that HP1 is required intrinsically to promote GSC self-renewal and progeny differentiation by directly stabilizing the transcripts of key genes involved in GSCs maintenance.

## Introduction

Stem cells are undifferentiated cells defined by their unique capacity to maintain self-renewing potential at every cell division, while producing differentiating daughter cells to ensure the correct development and maintain tissues homeostasis^1-3^. A better understanding of stem cells biology will not only reveal the crucial molecular mechanisms that control the formation and maintenance of tissues, but will also influence stem cell-based therapies in regenerative medicine^2,4,5^ and cancer treatments^6^.

In view of this, deepening the molecular mechanisms that control the fine balance between stem cell self-renewal and differentiation represents one of the fundamental goals of stem cell biology. This balance often depends on the coordinated regulation of complex transcriptional and post-transcriptional hierarchies.

The best way to investigate the molecular basis of stem cell regulation involves *in vivo* approaches, in the whole organism, since the removal of stem cells from the contexts of their "niches", in tissue cultures, could irreversibly change their properties^7^. In this context, the *Drosophila* ovarian germline stem cells (GSCs) represent an excellent and attractive model system to study the molecular basis of adult stem cell behavior and regulation^8-11^.

The *Drosophila* ovary is composed of about 20 functional units called ovarioles^12^. The most anterior part of the ovarioles consist of a germarium, a structure containing two or three asymmetrically dividing germline stem cells each of which produce another self-renewing GSC that remains anchored to the stromal somatic cap cells and a cystoblast (CB) committed to differentiate to sustain the later stages of the oogenesis.

The CB undergoes four synchronous divisions with incomplete cytokinesis to produce a 16-cell germ line cyst^12,13^ and steadily moves in a posterior direction through the germarium. Of these, one cell will differentiate into an oocyte, while the remaining cells will become polyploidy nurse cells^14^. The 16 cells cyst becomes surrounded by a monolayer of follicle cells and buds off from the posterior germarium to form an egg chamber^15,16^ which ultimately gives rise to a single mature oocyte ready for fertilization.

The activity of GSCs is controlled by extrinsic and intrinsic signaling pathways that finely regulate the balance between stem cell self-renewal and differentiation through the coordination of complex transcriptional and post-transcriptional hierarchies.

Decapentaplegic (Dpp) and Glass bottom boat (Gbb) are produced from the somatic niche and activate bone morphogenetic protein (BMP) signaling in the GSC to directly repress the Bam-dependent differentiation pathway and to maintain GSC identity^17-20^. Besides extrinsic mechanisms, stem cell intrinsic programs are crucial to control the binary germ line cell fate in *Drosophila*.

Nanos and Pumilio are intrinsic factors essential to maintain stem cell identity^21-23^. They are key components of an evolutionarily conserved translational repressor complex^24-28^ that bind to specific recognition sequences in the 3’untranslated regions (3’UTRs) of differenziating mRNAs to repress their translation^24,29,30^.

Other intrinsic factors necessary for GSC maintenance include components of the microRNA (miRNA) silencing machinery, indicating a central role for miRNA-dependent gene silencing in GSC identity^31-34^. Additionally, many genes involved in piRNA pathway appear to be crucial for proper GSC lineage development in *Drosophila*^22,35-38^.

A fast-growing body of experimental data provide strong evidences that also epigenetic mechanisms involving chromatin architecture and histone modification are equally important for the regulation of GSC maintenance and differentiation in *Drosophila*^39-43^. For example, the chromatin remodeling factor Iswi and the putative transcription factor Stonewall are intrinsically required for GSC maintenance^39-43^. The H3K4 demethylase Lsd1 controls non-autonomously the germ cell differentiation presumably through repressing *dpp* expression^41^. Moreover, other interesting studies show that the histone H2B ubiquitin protease Scrawny (Scny)^40^ and the histone H3K9 trimethylase Eggless (Egg) are required for maintaining self-renewal of GSC^42^.

Although different experimental evidence confirms the relevance of epigenetic regulatory programs in the GSC regulation, a complete picture of such mechanisms is still far to be resolved.

Heterochromatin protein 1 (HP1) is an evolutionarily conserved multifunctional epigenetic adaptor that is involved in heterochromatin formation and epigenetic gene silencing in different species including humans^44-46^. In addition to its role in heterochromatin structural organization, emerging evidence in *Drosophila* and mammals has highlighted the importance of HP1 in telomere capping, telomere length homeostasis^47,48^ and, more surprisingly, in positive regulation of gene expression^49-54^.

A recent study showed that HP1 and Su(var)3-9 are both necessary for GSC maintenance and that HP1 is sufficient for GSC self-renewal in *Drosophila* testis^55^. It has also been demonstrated that planarian HP1, induced upon injury, is able to promote regenerative proliferation of adult stem cells^56^. In mice, loss of HP1 gamma significantly reduces the number of primordial germ cells (PGCs) by regulating their cell cycle progression^57^. Moreover, HP1 gamma is essential for male germ cell survival and spermatogenesis^58^. Recently, a large-scale RNAi screen in *Drosophila* female germline stem cells identified HP1 as potentially involved in oogenesis^59^ even though the precise molecular mechanisms by which it exerts its function still remain elusive and need to be defined.

Here, we report our experiments showing an important function for *Drosophila* HP1 in female gametogenesis. In this study, we establish that HP1 is necessary for *Drosophila* oogenesis and is required cell autonomously to control the fine balance between stem cell self-renewal and differentiation. Finally, we show that HP1 exerts its functions, positively regulating the stability of key mRNAs involved in the control of female germ line stem cells development.

## Results and Discussion

### Functional inactivation of HP1 by *in vivo* RNA interference (RNAi) causes severe germ line defects that result in agametic ovarioles and female sterility

HP1 is a protein constitutively expressed in almost all larval and adult tissues with highest enrichment in adult ovaries (flybase.org). Immunostaining experiments performed by a specific anti-HP1 antibody on wild type ovaries, showed that HP1 localizes in the nucleus of both somatic and germline cells, from the anterior tip of the germarium (GSCs and CBs) until late stages of oogenesis (Fig. 1a). Specifically, HP1 immunosignals were mainly detected in dense pericentric heterochromatic foci in all germarium and developing egg chamber cells; HP1 also accumulated in the germinal vesicle and on the karyosome of the oocyte (Fig. 1a). HP1 was particularly enriched within and next to heterochromatic regions also in larval and pupal gonads (Supplementary Fig. S1).

**Figure 1.**
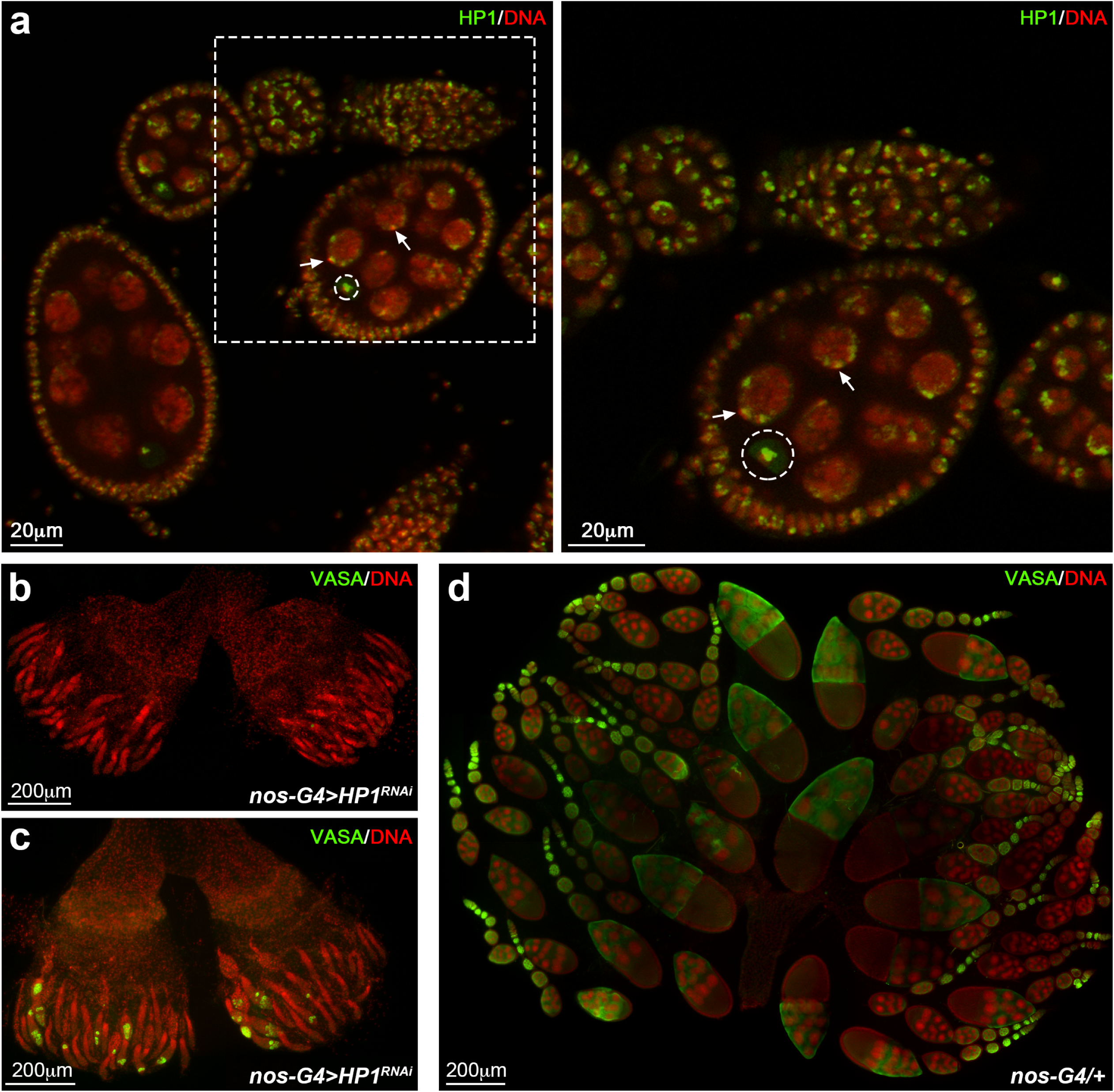
HP1 is required for correct ovarian development. **(a)** Wild type ovariole stained for DNA (red) and HP1 (green). Arrows indicate HP1 concentrated at domains of constitutive heterochromatin. Karyosome is identified by a white dashed circle. Dashed box is magnified in the right panel. **(b, c)** HP1 depleted ovaries stained for Vasa (green) and DNA (red). All ovarioles show an altered ovarian morphology, consisting in germaria completely empty **(b)** or germaria with only few germ cells **(c)**. **(d)** Developing wild-type ovaries obtained from newly eclosed females stained for Vasa (green) and DNA (red).

Since homozygous HP1 mutants die at third instar larvae, to investigate *in vivo* the function of HP1 in adult female germline, we took advantage of the Gal4-UAS binary system^60^. We performed tissue-specific HP1 knockdown by independently crossing two different transgenic lines carrying HP1 short small hairpin RNAs (shRNA)^61^ under the control of Gal4-responsive UAS promoter, with nanos-Gal4-NGT (hereafter referred as nos-Gal4) that provides a robust and uniform Gal4 expression in the germarium^62^. We found that the functional inactivation of HP1 in the F1 female progeny resulted in complete sterility thus suggesting an essential role for HP1 in female gametogenesis.

In order to further investigate the molecular basis underlying this female sterility, ovaries from *nos-Gal4>HP1^RNAi^* females ranging from 1-to 15-day-old, were dissected and immunostained with a specific antibody against Vasa, a DEAD-box RNA helicase which is a well-characterized marker of germ cells lineage in insects and vertebrates^63,64^. We found that knocking down HP1 upon nos-Gal4 driver expression (Supplementary Fig. S2), resulted in ovaries that were completely agametic (Fig. 1b, c) as compared to control ovaries (Fig. 1d); 86% of HP1 depleted germaria from 0-to 1-day-old females were completely devoid of germ cells (Fig. 1b) whereas 14% contained only a few germ cells at the tip of the ovariole (less than 10 per germarium) and one or two abnormal egg chambers (Fig. 1c) (n = 250 ovaries). From 5- to 15-day-old females, all the HP1 depleted ovaries exhibited a typical germline less morphology confirmed by the total absence of Vasa-positive cells (data not shown).

These findings strongly suggest for HP1 a specific and crucial role in germ line stem cell maintenance and differentiation; we could not, however, completely exclude a general role for HP1 in cell viability.

To discriminate between these possibilities, we knocked down HP1 with a maternal tubulin (Mat) Gal4 that induces transgenic expression of short hairpin RNAs against HP1 outside the germarium, starting in stage 2^65^ (Supplementary Fig. S3a, b). We found that HP1 knockdown females were fertile and showed no obvious oogenesis defects (Supplementary Fig. S3c) thus suggesting for HP1 an essential and cell autonomous function in early oogenesis and not a general requirement for cell survival.

### HP1 is required during multiple processes in early oogenesis

Germ cell-specific knockdown of HP1 causes almost complete loss of germ cells before adulthood. In order to determine the phenocritical period for HP1 requirement during normal oogenesis, we cytologically examined larval and pupal HP1 depleted ovaries following the germ cells fate, starting from early stages of germ-cell development to adulthood (Fig. 2 and Supplementary Fig. S4) Vasa staining analysis showed that larval ovaries from *nos-Gal4>HP1^RNAi^* females displayed a normal cellular organization as compared to control; consistent with this the total number of PGCs resulted unaffected (*nos-G4/+*, 107.5 ± 8.5; *nos-G4>HP1^RNAi^*, 106.6 ±7.0) (Fig. 2a).

**Figure 2.**
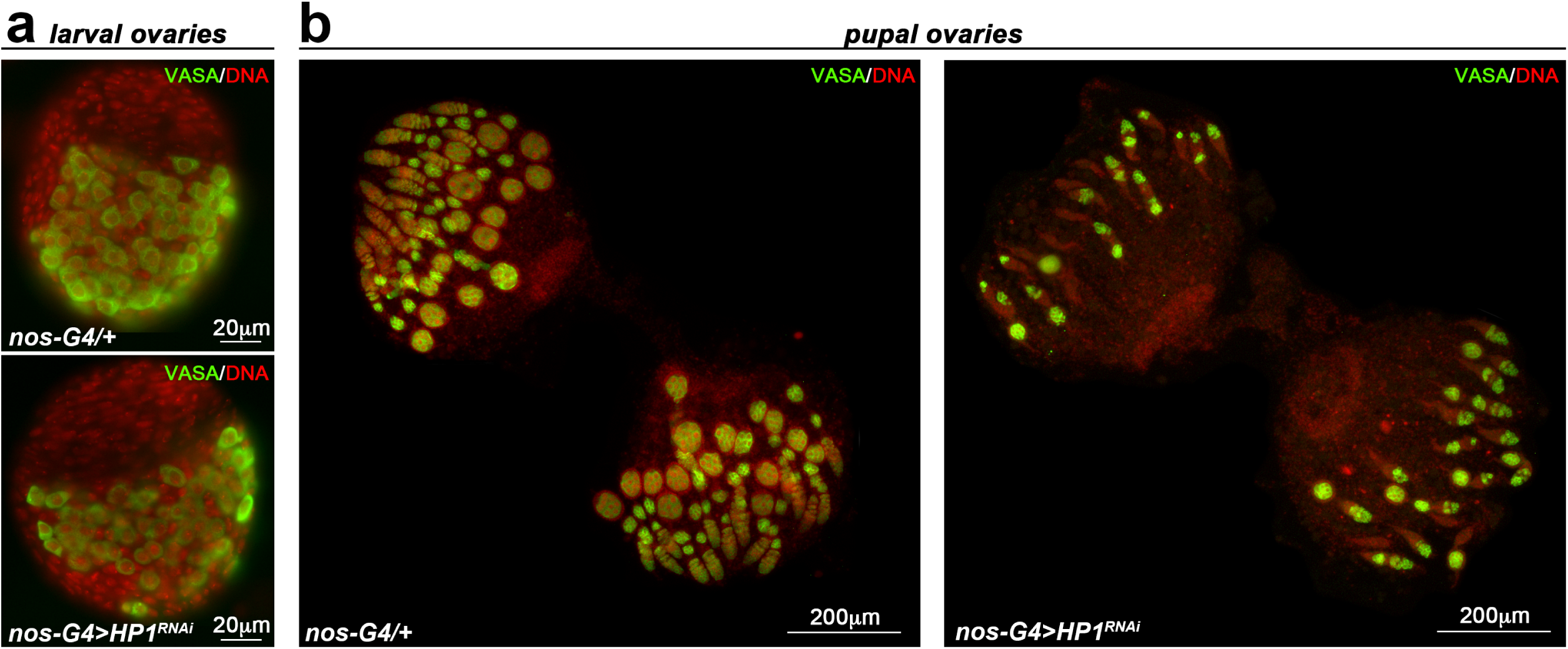
HP1 is required during the earliest stages of oogenesis at the transitional period of pupal stage. **(a)** Developing ovaries obtained from female wandering third-instar larvae stained for Vasa (green) and DNA (red). **(b)** Developing ovaries obtained from 72-96 h old pupae stained for Vasa (green) and DNA (red).

On the contrary, HP1 depleted pupal ovaries were almost completely devoid of differentiated egg chambers when compared to the control pupal gonads (Fig. 2b).

Taken together, these findings suggest that HP1 is required during the earliest stages of oogenesis at the larval/pupal transition when GSCs are established^66^.

In order to gain a more complete understanding of the altered phenotypes observed in pupal ovaries and to better investigate how HP1 regulates the behavior of germ cells, we performed an accurate cytological analysis on *nos-Gal4>HP1^RNAi^* pupal ovaries. We performed double-immunostaining experiments with antibodies against Vasa and α-Spectrin; α-Spectrin is a cytoskeletal protein that specifically labels spectrosomes and fusomes and can be used to trace the germline differentiation. Spectrosomes are spherical and mark GSCs and cystoblasts, whereas fusomes are branched and mark 2, 4, 8, and 16-cell cysts^67^.

The results of this cytological analysis (152 ovarioles scored) demonstrated that HP1 depleted pupal ovaries exhibited several remarkable and complex phenotypes including: empty germaria (27%, n=41 ovarioles), germaria with germ cells containing spectrosomes only (33%, n=51 ovarioles), germaria with germ cells containing both spectrosomes and fusomes carrying a single developing cyst connected by an incompletely branched fusome (26%, n=39 ovarioles) and germaria with few germ cells containing only fusomes (14%, n=21 ovarioles) (Fig. 3a, b).

**Figure 3.**
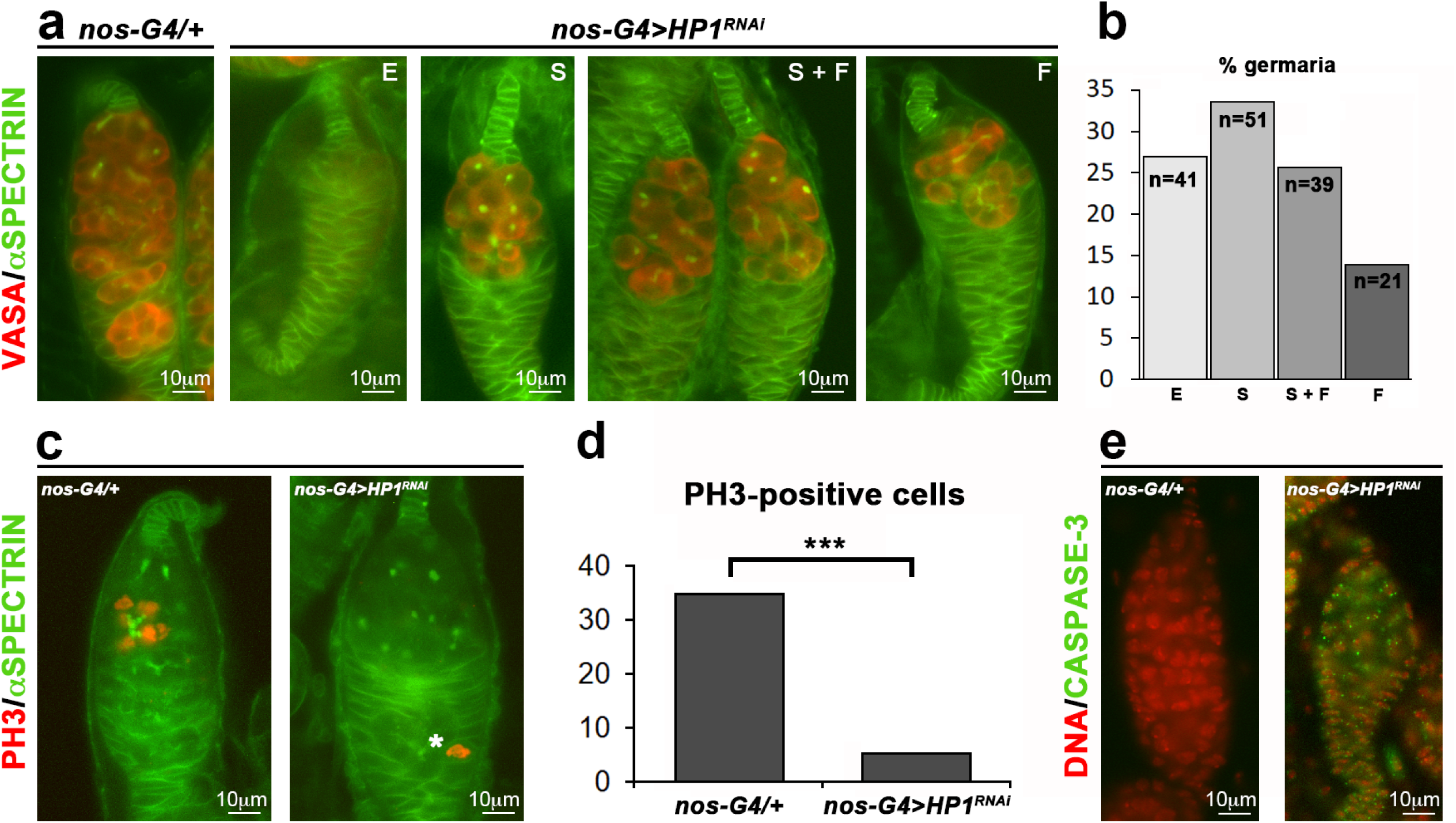
Loss of HP1 Causes a Complex GSC Phenotype. **(a)** Representative images of each phenotypic class obtained from pupal HP1-depleted germaria (*nos-G4>HP1^RNAi^*) stained for Vasa (red) and α-Spectrin (green). **(b)** Quantification of the prevalence of each phenotypic class in HP1 depleted pupal ovaries; the total number of germaria scored is shown within each bar. E, empty germaria; S, germaria with spectrosomes only; S+F, germaria with germ cells containing both spectrosomes and fusomes; F, germaria with only fusome-containing germ cells. **(c)** Double-staining immunofluorescence on control (*nos-G4/+*) and HP1 depleted (*nos-G4>HP1^RNAi^*) pupal ovaries for α-Spectrin (green) and PH3 (red). The white asterisk indicates dividing follicle stem cell (FSC). **(d)** Quantification of PH3-positive cystoblast in HP1 knockdown pupal ovaries. Statistical significance was determined by Fisher’s exact test (\*\*\**p* <0.001). **(e)** Immunofluorescence on control (*nos-G4/+*) and HP1 depleted (*nos-G4>HP1^RNAi^*) pupal ovaries for cleaved Caspase-3 (green) and DNA (red).

These complex phenotypic defects suggest for HP1 a functional role in regulating the germline stem cell (GSC) maintenance.

We asked whether the low number of germ cells in HP1 depleted ovarioles could be related to defects in the division rate of ovarian stem cells and their progeny. These defects might contribute to germ line cells loss over time. In order to verify the capacity of germ cells to undergo mitotic divisions, we immunostained wild type and HP1 knockdown ovaries with a specific antibody to phosphorylated H3S10 (phospho-H3, PH3) to detect germline cells undergoing mitosis at a given time (Fig. 3c). In HP1 depleted ovaries we observed an almost complete loss of PH3 positive nuclei (5%, n=56 ovarioles) respect to the control ovaries (35%, n=46 ovarioles) (Fig. 3d); this result establishes that the functional inactivation of HP1 severely impairs the correct germ cells division. We also assessed apoptosis by using anti-cleaved Caspase-3 antibody that is a proven marker for cells that are dying. The results clearly indicated that the few remaining germline cells detected in HP1 depleted ovaries are strongly stained with cleaved Caspase-3 suggesting that the germ cells that fail to properly divide die prematurely (Fig. 3e).

### HP1 promotes germ cell differentiation by post-transcriptionally regulating *bam* expression and function

Germline division defects are often associated to an altered differentiation program. Previous studies demonstrated that Bag of Marbles (Bam) protein is necessary and sufficient for promoting GSC and cystoblasts differentiation, since *bam* mutations completely block germ cell differentiation (causing GSC hyperplasia), whereas *bam* ectopic expression in GSCs results in their complete and precocious differentiation^18,68,69^.

To determine whether the phenotypic defects observed in HP1 depleted pupal ovaries could be related to *bam* repression, we firstly evaluated, by quantitative real-time PCR (qRT-PCR), the expression of *bam* gene in HP1 knockdown pupal ovaries. We found that ovaries lacking HP1 exhibited a significant reduction of *bam* transcript levels (close to about 80%) as compared to control ovaries (Fig. 4a).

**Figure 4.**
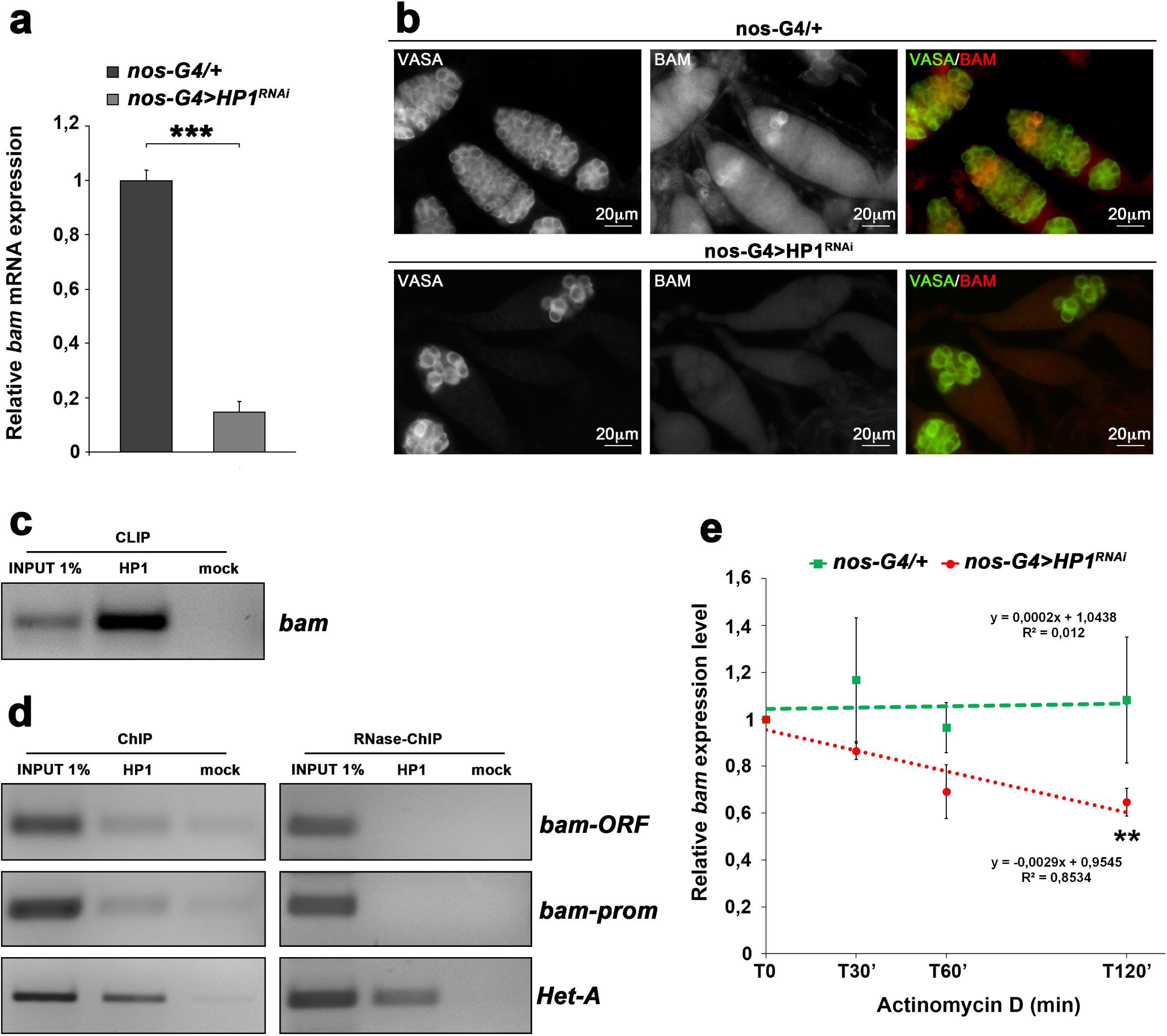
HP1 regulates *bam* mRNAs in a post-transcriptional manner. **(a)** qRT-PCR analysis showing that HP1 depleted pupal ovaries express significantly less *bam* transcript respect to the control. Fold-changes in RNA levels relative to the control were normalized to *rp49* levels. Error bars indicate ±SEM from three biological replicates (\*\*\**p* < 0.001). **(b)** Double immunofluorescence on control (*nos-G4/+*) and HP1 depleted (*nos-G4>HP1^RNAi^*) pupal ovaries for Vasa and Bam. **(c)** RT-PCR analysis of RNAs immunoprecipitated with α-HP1 (HP1 CLIP sample) in newly eclosed females ovaries. The PCR data shown here are representative of three independent CLIP experiments. The full-length versions of the cropped gels are reported in Supplementary Fig. S7a. **(d)** Chromatin immunoprecipitation (ChIP) analysis of HP1 occupancy at the *bam* promoter region (bam Silencer Element) and coding sequence in newly eclosed female ovaries. The RNase sensitivity of this association was tested by pre-treating the extract with a combination of RNase A and RNase T1 (right panel). Het-A was used as a positive control to check whether the ChIP experiments were working. PCR reactions were carried out on 1% input DNA. The PCR data shown here are representative of three independent ChIP experiments. The full-length versions of the cropped gels are reported in Supplementary Fig. S7b, c. **(e)** qRT-PCR analysis of *bam* mRNA transcript at different times after blockage of transcription by Actinomycin D treatment. The green line and the red line indicate the *bam* transcript amount respectively in the control (*nos-G4/+*) and HP1-depleted (*nos-G4>HP1^RNAi^*) ovaries from 1-day-old females. Total RNA was isolated at the indicated times (0, 30 min, 60 min and 120 min). The values shown are averages ±SEM of three biological replicates. The dashed lines represent the best fit regression of all data point and the slopes are shown on the graph. For each genotype, all data point vs T0 was statistically evaluated by one-sample t-test (\*\**p* < 0.01).

Consistent with the down regulation of *bam* mRNAs, we also observed a drastic diminution of Bam protein by immunostaining with a specific monoclonal antibody against Bam (Fig. 4b). In wild type ovaries, Bam protein was detected, as expected, in cystoblasts and early developing cysts (2-, 4-, and 8-cell cysts) whereas in HP1 mutant ovaries Bam protein was almost undetectable (Fig. 4b). Altogether, these data strongly suggest that HP1 blocks Bam driving germ cell differentiation. Previously we have demonstrated that in *Drosophila* HP1 takes part in positive regulation of gene expression by stabilizing RNA transcripts and protecting them against premature and rapid degradation^53^; in particular, we found that HP1 is able to directly bind the transcripts of more than one hundred euchromatic genes in *Drosophila* and physically interacts with DDP1^70^, HRB87F^71^ and PEP^72^, which belong to different classes of heterogeneous nuclear ribonucleoproteins (hnRNPs) that are known to be involved in RNA packaging, stability and processing. Moreover, in our previous work we also demonstrated that HP1 is cotranscriptionally recruited on nascent transcripts through its chromodomain^49,53^.

In order to verify if HP1 was directly involved in post-transcriptional regulation of *bam* gene by binding *in vivo* its mRNA, we performed HP1 CLIP (UV cross-linking and immunoprecipitation) experiments^73,74^ on whole adult ovaries dissected from 0-to 1-day-old-wild type females. The results of RT-PCR from HP1 CLIP experiments clearly showed that *bam* transcripts were significantly enriched in the CLIP sample when compared to the mock control sample (Fig. 4c) and demonstrated that HP1 is able to specifically bind *bam* transcripts *in vivo*. In order to further investigate whether HP1 is cotranscriptionally recruited on *bam* nascent transcripts, we performed ChIP experiments on cross-linked chromatin purified from 0- to 1-day-old wild-type ovaries. To evaluate the presence of *bam* sequences among the immunoprecipitated DNA, a PCR analysis was performed with specific primer pairs covering both the promoter and the coding regions of *bam* gene.

The results of ChIP assays demonstrated that HP1 is clearly associated to *bam* gene (Fig. 4d). To completely exclude any direct role for HP1 on *bam* transcriptional control and to confirm that HP1 binding on *bam* gene was exclusively mediated by the presence of *bam* nascent transcripts, ChIP experiments were repeated in presence of RNaseA/T1 mix that specifically degrades single stranded RNA (ssRNA). The RNase-ChIP results demonstrated that chromatin RNase treatment prior to immunoprecipitation completely remove HP1 from *bam* gene thus confirming that the recruitment of HP1 on *bam* gene is clearly RNA-dependent (Fig. 4d). RNase treatment did not affect, as expected, the HP1 occupancy over Het-A telomeric retrotransposon (Fig. 4d) since, at the telomeres, HP1 is capable to directly bind HeT-A sequences through its hinge domain^48^. To determine the stability of *bam* transcripts, we analyzed, by qRT-PCR, RNA samples purified from wild type and HP1 knockdown ovaries treated with Actinomycin D to inhibit transcription and *de novo* RNA synthesis. Previous analysis showed that a 30 min treatment was sufficient to inhibit transcription in the ovaries^75^. As shown in Figure 4e, in HP1 lacking ovaries we observed a strong and rapid decay rate of *bam* transcript when compared to the control (Fig. 4e).

These observations strongly suggest that HP1 may regulate *bam* mRNAs in a post-transcriptional manner.

To confirm our findings and to verify if HP1 can effectively control germ cells differentiation in a *bam*-dependent manner, we overexpressed *bam* from a heat shock inducible transgene carrying the full-length bam cDNA^68^ in the HP1 knockdown germ cells. To assess the effectiveness of hs-bam transgene expression we analyzed bam mRNA and protein in HP1 depleted ovaries with or without heat-shock (Supplementary Fig. S5).

*Nos-Gal4 /UAS-HP1^RNAi^; P[hs-bam]/+* and *nos-Gal4 /HP1^RNAi^; +/+* females were heat-shocked at pupal stage (96 hours) at 37 °C for 1 hour and, 24 hours after heat shock (HS) treatment, adult ovaries were dissected and stained with anti-Vasa antibody (Fig. 5a, b). As showed in Figure 5b, heat-shock induced *bam* can only partially rescue the phenotypic defects induced by HP1 knockdown since its forced expression under control of the heat shock promoter generates only few normally developed egg chambers (see Fig. 5c for quantification of ovarioles containing developing egg chambers in heat shocked HP1 depleted females carrying the *P[hs-bam]* transgene). This finding suggests that oogenesis defects observed in HP1 depleted ovaries may be only partially imputable to defective differentiation mechanisms.

**Figure 5.**
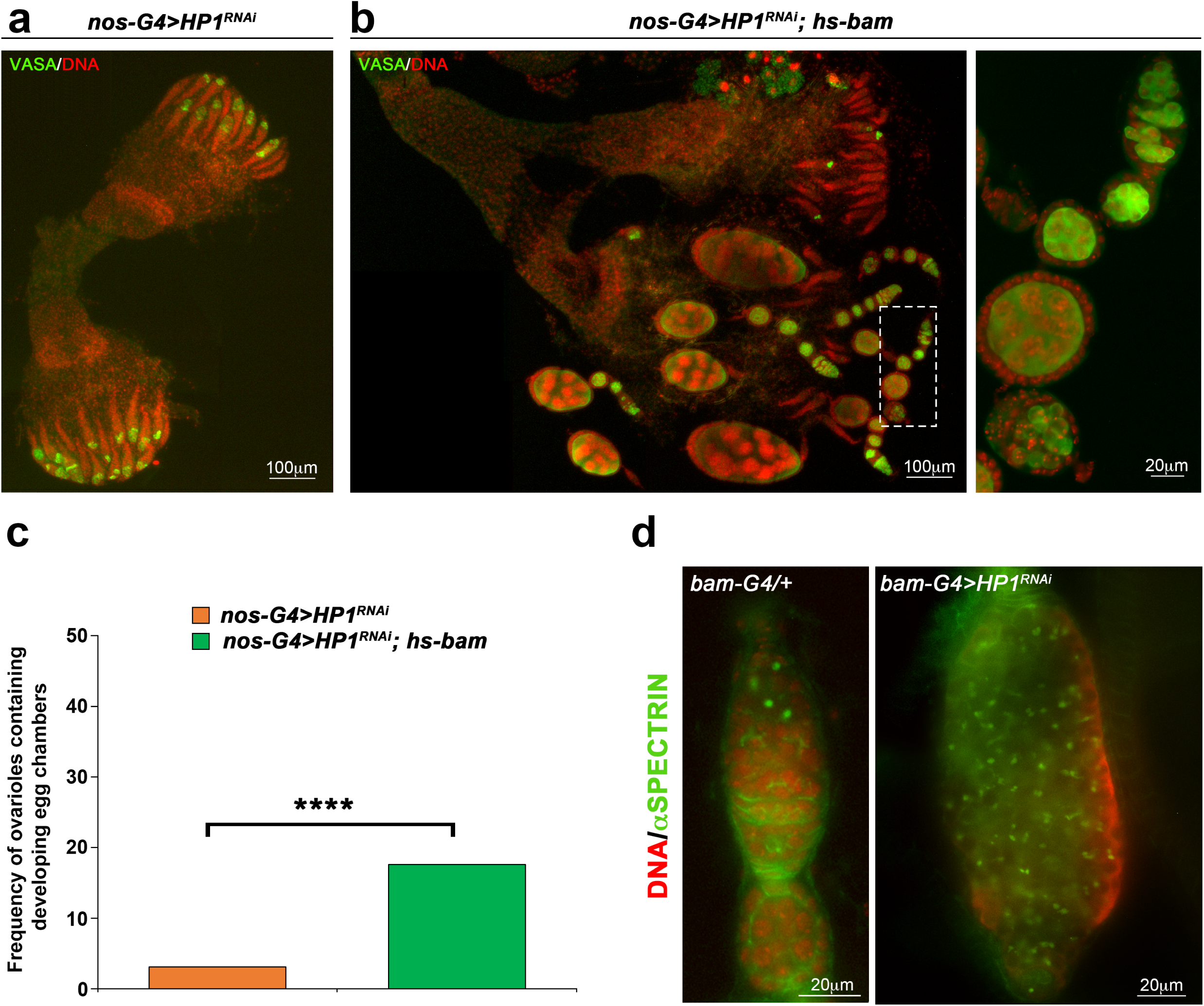
Heat-shock induced *bam* can only partially rescue the phenotypic defects induced by HP1 knockdown. **(a,b)** Staining for Vasa (green) and DNA (red) on whole mounts adult ovaries from *nos-G4>HP1^RNAi^* **(a)** and *nos-G4>HP1^RNAi^; hs-bam* **(b)** females. **(b)** The dashed white box in the left panel is magnified in the right panel. **(c)** Quantification of ovarioles containing developing egg chambers in heat shocked HP1 depleted females carrying or not the *P[hs-bam]* transgene (454 and 784 ovarioles scored for *nos-G4>HP1^RNAi^* and *nos-G4>HP1^RNAi^; hs-bam,* respectively). Statistical significance was determined by Fisher's exact test (\*\*\*\**p* < 0.0001). **(d)** Staining for Spectrin (green) and DAPI (red) on germaria obtained from *bam-G4>HP1^RNAi^* females.

It is well known that loss of *bam* blocks germ cell differentiation resulting in GSC hyperplasia^68^, a characteristic phenotype that we never observed in HP1 depleted ovaries by nos-Gal4.

Altogether, these findings strongly suggest that the complex phenotypic defects arising from HP1 knockdown in the female germline are only partially dependent on *bam* repression and are probably due to a duplex coordinated control operated by HP1 in both GSCs self-renewal and differentiation. In order to verify this hypothesis we inactivated HP1 only in Bam-expressing germline cells by using P{bam promoter-Gal4:VP16}^17^ that drives the expression of shHP1 only in the dividing cystoblast and cystocytes but not in GSCs where the function of HP1 protein remains completely wild-type. In this case, we observed the classical ovarian tumor phenotype (Fig. 5d) albeit at very low frequency (less than 1%) due to the low effectiveness of bam-Gal4 driver in knocking down HP1 protein (Supplementary Fig. S6).

### HP1 controls GSCs self-renewal by post-transcriptional regulation of stemness genes

Consistent with the conclusion stated above, we wondered if HP1 was able to post-transcriptionally regulate also key stemness genes. First, we analyze by qRT-PCR the expression profiles of some important genes that are intrinsically involved in GSCs self-renewal by repressing Bam differentiation pathways^76-78^.

We found that some of them as *nos*, *cup*, *piwi* and *vasa* were significantly down regulated in HP1 knockdown pupal ovaries respect to the control (Fig. 6a). These results allowed us to hypothesize that also *nos*, *cup*, *piwi* and *vasa* genes might be post-transcriptionally regulated by HP1. So we dissected ovaries from 0- to 1-day-old wild type females to repeat both CLIP and ChIP experiments. CLIP-PCR analysis, clearly showed that *nos*, *cup* and *piwi* RNAs were significantly enriched in the IP sample respect to the mock control sample whereas *vasa* mRNA did not (Fig. 6b). These genes resulted strongly enriched also in ChIP IP sample but not in RNAse-ChIP IP sample (Fig. 6c) indicating that their RNAs are co-transcriptionally bound by HP1. To determine the mRNA decay of these genes, we repeated the Actinomycin D treatment that allowed us to conclude that HP1 is able to stabilize *nos*, *cup* and *piwi* mRNAs (Fig. 6d).

**Figure 6.**
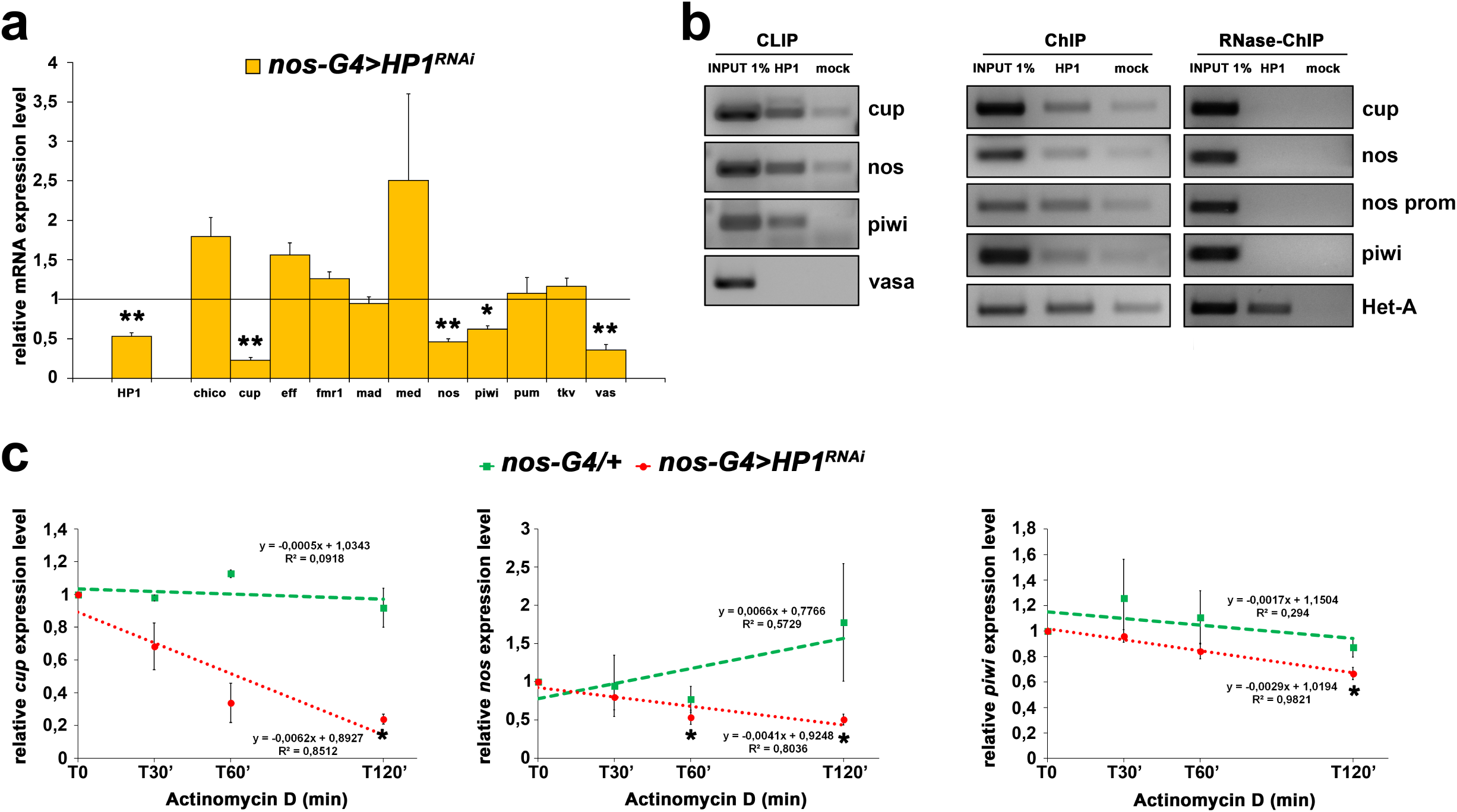
HP1 is required for GSC self-renewal. **(a)** Bar graph showing relative quantification of mRNA encoding GSCs key factors in HP1-depleted or control pupal ovaries. Error bars indicate ±SEM from three biological replicates (\*\**p* < 0.01, \**p* < 0.05). **(b)** RT-PCR analysis of RNAs immunoprecipitated with α-HP1 (HP1 CLIP sample) in ovarian extract from newly eclosed females. The PCR data shown here are representative of three independent CLIP experiments. The full-length versions of the cropped gels are reported in Supplementary Fig. S7a. **(c)** Chromatin immunoprecipitation (ChIP) analysis of HP1 occupancy at *cup*, *nos, piwi* and *vasa* genes in newly eclosed female ovaries. The RNase sensitivity of this association was tested by pretreating the extract with a combination of RNase A and RNase T1 (right panel). Het-A was used as a positive control. PCR reactions were carried out on 1% input DNA. The PCR data shown here is representative of three independent ChIP experiments. The full-length versions of the cropped gels are reported in Supplementary Fig. S7b, c. **(d)** Time course degradation assay of *cup, nos*, and *piwi* RNAs measured by qRT-PCR in control (*nos-G4/+*) or HP1 depleted (*nos-G4>HP1^RNAi^*) ovaries from 1-day-old females. Total RNA was isolated at the indicated times (0, 30 min, 60 min and 120 min). The values shown are averages ±SEM of three biological replicates. The dashed lines represent the best fit regression of all data point and the slopes are shown on the graph. For each genotype, all data point vs T0 was statistically evaluated by one-sample t-test (\**p* < 0.05).

Altogether, these data strongly indicate that HP1 is intrinsically required for post-transcriptional regulation of *Drosophila* GSC maintenance. Our findings suggest that HP1 exerts its function through the formation of an HP1-containing hnRNP nuclear complex that protects and stabilizes key mRNAs involved in the control of GSC homeostasis and behavior. Intriguingly there are different experimental evidences demonstrating that also mutations in genes coding for the HP1-interacting hnRNPs DDP1, HRB87F and PEP induce female-sterile phenotypes^79-81^. For example, the hypomorphic insertional allele of DDP1 (*Dp1^15.1^*) causes complete sterility in females but not in males. Homozygous *Dp1^15.1^* females show abnormal ovaries with ovarioles undeveloped, egg chambers often fused and containing an irregular number of cells^79^; finally, also PEP and HRB87F are essential for normal gonadal development and female fecundity^80,81^.

In conclusion, the above results demonstrate, for the first time, an essential role for HP1 in post-transcriptional regulation of GSC maintenance and certainly add a new dimension to our understanding of HP1 targeting and functions in epigenetic regulation of GSC behavior.

## Methods

### *Drosophila* Strains

All flies were raised at 24 °C on standard cornmeal-sucrose-yeast-agar medium.

For a detailed list of all stocks used in this study, see Supplementary Methods.

### Immunofluorescent staining of larval, pupal and adult whole-mount ovaries

Pupal and adult ovaries were stained according to Grieder^82^.

Larval ovaries were dissected, fixed, and immunostained as described previously by Pisano^83^.

Further details can be found in Supplementary Methods.

### Western Blot analysis

Protein extracts fractionated by 10% SDS-PAGE and electroblotted onto Immobilion-P polyvinyldifluoride membranes (Bio-rad) were probed with antibodies against HP1 (1:500, 9A9 monoclonal mouse), α-Tubulin (mouse 1:2000, Sigma). Proteins of interest were detected with HRP-conjugated goat anti-mouse or anti-rabbit IgG antibody (1:10000, Santa Cruz) and visualized with the ECL Western blotting substrate (GE Healthcare), according to the provided protocol. The chemiluminescence detection was performed on the ChemiDoc XRS+ System (Bio-rad) and analyzed using the included ImageLab software.

### Cross-linking immunoprecipitation (CLIP) assay

CLIP assay was performed as previously reported^84^ with some modifications. Approximately 20 mg ovaries from 0- to 1-day-old wild-type females were UV crosslinked (3×2000 µJ/cm^2^), homogenized on ice in 1 mL RCB buffer (50 mM HEPES pH 7.4, 200 mM NaCl, 2.5 mM MgCl_2_, 0.1% Triton X-100, 250 mM sucrose, 1 mM DTT, 1× EDTA-free Complete Protease Inhibitors, 1 mM PMSF) supplemented with 300 U RNAseOUT (Invitrogen) and placed on ice for 30 min. The homogenate was sonicated on ice, at 80% power, five times in 20 s bursts with a 60 s rest in between using the Hielscher Ultrasonic Processor UP100H (100W, 30kHz) and centrifuged (16000xg for 5 min at 4 °C). Soluble extract was precleared with 20 µl Protein-G dynabeads (Invitrogen) for 20 min at 4 °C. After removal of samples for immunoblotting and quantitation of RNA input (1%), HP1 was immunoprecipitated with anti-HP1 9A9 antibody from 450 µl precleared extract by incubation for 4 h with 50 µl Protein-G dynabeads. Immunoprecipitates were washed 4 times with RCB. To elute the immunoprecipitated RNAs, the pelleted beads were boiled in 100 µL of UltraPure DEPC-treated Water for 5 min. 900 µL Qiazol Reagent was added to the supernatant recovered for RNA preparation. The RNA purified was used as a template to synthesize cDNA using oligo dT, random hexamers and SuperScript reverse transcriptase III (Invitrogen) according to the manufacturer's protocol.

### Chromatin immunoprecipitation assay

Chromatin immunoprecipitation was performed according to the method described by Menet^85^ with minor modifications. Approximately 20 mg ovaries from 0- to 1-day-old wild-type females were homogenized in 1 mL of NEB buffer (10 mM HEPES-Na at pH 8.0, 10 mM NaCl, 0.1 mM EGTA-Na at pH 8, 0.5 mM EDTA-Na at pH 8, 1 mM DTT, 0.5% NP-40, 0.5 mM Spermidine, 0.15 mM Spermine, 1× EDTA-free Complete Protease Inhibitors) with a Polytron homogenizer (Kinematica Swizerland) with a PT300 tip for 1 min (at 3000 rpm). The homogenate was transferred to a pre-chilled glass dounce (Wheaton) and 15 full strokes were applied with a tight pestle. Free nuclei were then centrifuged at 6000xg for 10 min at 4 °C. The nuclei-containing pellets were resuspended in 1 mL of NEB and centrifuged at 20000xg for 20 min on sucrose gradient (0.65 mL of 1.6 M sucrose in NEB, 0.35 mL of 0.8 M sucrose in NEB). The pellet was resuspended in 1 mL of NEB and formaldehyde to a final concentration of 1%. Nuclei were cross-linked for 10 min at room temperature and quenched by adding 1/10 vol of 1.375 M glycine. The nuclei were collected by centrifugation at 6000xg for 5 min. Nuclei were washed twice in 1 mL of NEB and resuspended in 1 mL of Lysis Buffer (15 mM HEPES-Na at pH 7.6, 140 mM NaCl, 0.5 mM EGTA, 1 mM EDTA at pH 8, 1% Triton X-100, 0.5 mM DTT, 0.1% Na Deoxycholate, 0.1% SDS, 0.5% N-lauroylsarcosine and 1× EDTA-free Complete Protease Inhibitors). Nuclei were sonicated using a Hielscher Ultrasonic Processor UP100H (100W, 30kHz) six times for 20 s on and 1 min on ice. Sonicated nuclei were centrifuged at 13000xg for 4 min at 4 °C. The majority of sonicated chromatin was 500 to 1000 base pairs (bp) in length. For each immunoprecipitation, 15 µg of chromatin was incubated in the presence of 10 µg of HP1 9A9 monoclonal antibody (3 h at 4 °C in a rotating wheel). Then, 50 µl of dynabeads protein G (Invitrogen) was added and incubation was continued overnight at 4 °C. The supernatants were discarded and samples were washed twice in Lysis Buffer (each wash 15 min at 4 °C) and twice in TE Buffer (1 mM EDTA, 10 mM TrisHCl at pH 8). Chromatin was eluted from beads in two steps; first in 100 µl of Eluition Buffer 1 (10 mM EDTA, 1% SDS, 50mM TrisHCl at pH 8) at 65 °C for 15 min, followed by centrifugation and recovery of the supernatant. Beads material was re-extracted in 100 µl of TE + 0.67% SDS. The combined eluate (200 µl) was incubated overnight at 65 °C to reverse cross-links and treated by 50 µg/ml RNaseA for 15 min at 65 °C and by 500 µg/ml Proteinase K (Invitrogen) for 3 h at 65 °C. Samples were phenol–chloroform extracted and ethanol precipitated. DNA was resuspended in 25 µl of water. For maximising the molecular analyses with DNA immunoprecipitated, candidate genes were amplified in pairs through an optimized duplex-PCR protocol by using two different sets of primers having similar melting temperatures in a single reaction.

RNAse-Chromatin immunoprecipitation was performed essentially as described for ChIP but with an important modification: sheared chromatin was treated with RNAse mix (Roche) for 1 h at 37 °C before immunoprecipitation.

### Primers design and PCR amplification

All PCR specific primers (18–25 mers with a minimum GC content of 50% and average Tm of 60 °C) (listed in Supplementary Table S1) were designed using the Invitrogen OligoPerfect™ designer web tool and oligonucleotide sequences were screened using a BLAST search to confirm the specificity. PCR amplifications were performed with Platinum^®^ Taq DNA Polymerase Kit (Invitrogen) according to the manufacturer’s instructions.

The thermal profile for PCR amplification of CLIP samples was as follows: initial denaturation at 94 °C for 5 min, followed by 35 cycles of 94 °C for 30 s, 60 °C for 30 s, 72 °C for 30 s, and ending with a final extension at 72 °C for 7 min.

The thermal profile for duplex-PCR amplification of ChIP samples was as follows: initial denaturation at 94 °C for 5 min, followed by 28 cycles of 94 °C for 30 s, 60 °C for 30 s, 72 °C for 30 s, and ending with a final extension at 72 °C for 7 min. The PCR products were analyzed by 2% agarose gel electrophoresis.

### Total RNA extraction and qRT-PCR

RNA samples from ovaries were isolated by Qiazol reagent (Qiagen) according to the manufacturer’s instructions. The concentration and purity of RNAs were determined using NanoDrop 1000 Spectrophotometer (Thermo Scientific). 5 µg of total RNA was reverse transcribed using oligo dT and SuperScript Reverse Transcriptase III (Invitrogen) according to the manufacturer's protocol. The qPCR reactions were carried out with QuantiFast SYBR Green PCR Kit (Qiagen) according to manufacturer’s protocol. Relative abundance of the different transcripts was determined using the 2^−ΔΔCt^ method^86^ using *rp49* transcript as control. qRT-PCR experiments were performed in three independent biological replicates; all reactions were run in triplicates in 96-well plates over 40 cycles of 95 °C for 15 s and 60 °C for 60 s in a two-step thermal cycle preceded by an initiation step of 95 °C for 10 min. Melting-curve analysis was performed on each sample to control for nonspecific amplification and primer dimer formation. Primer sequences were listed in Supplementary Table S1. Statistical significance was determined by Mann-Whitney tests using GraphPad Prism Software. A *p* value ≤ 0.05 was considered statistically significant.

### Actinomycin D treatment

To assay for mRNA stability, ovaries dissected from 1-day-old females raised at lower temperature (18 °C) were treated with 20 μg/ml Actinomycin D in Schneider’s medium with constant rocking at room temperature for 30 min (T0, sufficient to inhibit transcription as described in Jao and Salic^87^; total RNA was extracted at T0 and then T30 min, T60 min, T120 min. mRNA levels for *bam*, *nos*, *piwi* and *cup* were analyzed by qRT-PCR.

## Supporting information

## Acknowledgements

We are grateful to S. Pimpinelli and C. Vincenzi for critical reading of the manuscript. We would like to thank S. Caristi for assistance with confocal image acquisition, B. Wakimoto, P. Dimitri and Developmental Studies Hybridoma Bank at the University of Iowa, for antibodies. We would like to thank also V. Palumbo, Bloomington *Drosophila* Stock Center and Vienna *Drosophila* RNAi Center for kindly providing fly stocks.

## Author contributions

A.M.C. and U.C. contributed to experimental design, performed, and analyzed all experiments; L.F. contributed to data analysis. L.P. designed, analyzed, supervised all experiments and wrote the manuscript. All authors reviewed the manuscript.

## Additional Information

### Competing Interests

The authors declare no competing interests.

## References

1 Lin, H. The stem-cell niche theory: lessons from flies. Nat Rev Genet 3, 931–940, doi:10.1038/nrg952 (2002).

2 Spradling, A., Drummond-Barbosa, D. & Kai, T. Stem cells find their niche. Nature 414, 98–104, doi:10.1038/35102160 (2001).

3 Morrison, S. J. & Spradling, A. C. Stem cells and niches: mechanisms that promote stem cell maintenance throughout life. Cell 132, 598–611, doi:10.1016/j.cell.2008.01.038 (2008).

4 Fuchs, E. J. & Whartenby, K. A. Hematopoietic stem cell transplant as a platform for tumor immunotherapy. Curr Opin Mol Ther 6, 48–53 (2004).

5 Mahla, R. S. Stem Cells Applications in Regenerative Medicine and Disease Therapeutics. Int J Cell Biol 2016, 6940283, doi:10.1155/2016/6940283 (2016).

6 Shah, K. Stem cell-based therapies for tumors in the brain: are we there yet? Neuro Oncol 18, 1066–1078, doi:10.1093/neuonc/now096 (2016).

7 Schofield, R. The relationship between the spleen colony-forming cell and the haemopoietic stem cell. Blood Cells 4, 7–25 (1978).

8 Fuller, M. T. & Spradling, A. C. Male and female Drosophila germline stem cells: two versions of immortality. Science 316, 402–404, doi:10.1126/science.1140861 (2007).

9 Spradling, A., Fuller, M. T., Braun, R. E. & Yoshida, S. Germline stem cells. Cold Spring Harb Perspect Biol 3, a002642, doi:10.1101/cshperspect.a002642 (2011).

10 Xie, T. et al. Interactions between stem cells and their niche in the Drosophila ovary. Cold Spring Harb Symp Quant Biol 73, 39–47, doi:10.1101/sqb.2008.73.014 (2008).

11 Singh, G. Drosophila’s contribution to stem cell research. F1000Res 4, 157, doi:10.12688/f1000research.6611.2 (2015).

12 Spradling, A. C. Germline cysts: communes that work. Cell 72, 649–651 (1993).

13 de Cuevas, M., Lilly, M. A. & Spradling, A. C. Germline cyst formation in Drosophila. Annu Rev Genet 31, 405–428, doi:10.1146/annurev.genet.31.1.405 (1997).

14 Cooley, L. & Theurkauf, W. E. Cytoskeletal functions during Drosophila oogenesis. Science 266, 590–596 (1994).

15 Forbes, A. J., Lin, H., Ingham, P. W. & Spradling, A. C. hedgehog is required for the proliferation and specification of ovarian somatic cells prior to egg chamber formation in Drosophila. Development 122, 1125–1135 (1996).

16 Margolis, J. & Spradling, A. Identification and behavior of epithelial stem cells in the Drosophila ovary. Development 121, 3797–3807 (1995).

17 Chen, D. & McKearin, D. M. A discrete transcriptional silencer in the bam gene determines asymmetric division of the Drosophila germline stem cell. Development 130, 1159–1170 (2003).

18 McKearin, D. M. & Spradling, A. C. bag-of-marbles: a Drosophila gene required to initiate both male and female gametogenesis. Genes Dev 4, 2242–2251 (1990).

19 Song, X. et al. Bmp signals from niche cells directly repress transcription of a differentiation-promoting gene, bag of marbles, in germline stem cells in the Drosophila ovary. Development 131, 1353–1364, doi:10.1242/dev.01026 (2004).

20 Xie, T. & Spradling, A. C. decapentaplegic is essential for the maintenance and division of germline stem cells in the Drosophila ovary. Cell 94, 251–260 (1998).

21 Forbes, A. & Lehmann, R. Nanos and Pumilio have critical roles in the development and function of Drosophila germline stem cells. Development 125, 679–690 (1998).

22 Lin, H. & Spradling, A. C. A novel group of pumilio mutations affects the asymmetric division of germline stem cells in the Drosophila ovary. Development 124, 2463–2476 (1997).

23 Wang, Z. & Lin, H. Nanos maintains germline stem cell self-renewal by preventing differentiation. Science 303, 2016–2019, doi:10.1126/science.1093983 (2004).

24 Sonoda, J. & Wharton, R. P. Recruitment of Nanos to hunchback mRNA by Pumilio. Genes Dev 13, 2704–2712 (1999).

25 Zamore, P. D., Williamson, J. R. & Lehmann, R. The Pumilio protein binds RNA through a conserved domain that defines a new class of RNA-binding proteins. RNA 3, 1421–1433 (1997).

26 Zhang, B. et al. A conserved RNA-binding protein that regulates sexual fates in the C. elegans hermaphrodite germ line. Nature 390, 477–484, doi:10.1038/37297 (1997).

27 Wickens, M., Bernstein, D. S., Kimble, J. & Parker, R. A PUF family portrait: 3’UTR regulation as a way of life. Trends Genet 18, 150–157 (2002).

28 Kimble, J. & Crittenden, S. L. Controls of germline stem cells, entry into meiosis, and the sperm/oocyte decision in Caenorhabditis elegans. Annu Rev Cell Dev Biol 23, 405–433, doi:10.1146/annurev.cellbio.23.090506.123326 (2007).

29 Murata, Y. & Wharton, R. P. Binding of pumilio to maternal hunchback mRNA is required for posterior patterning in Drosophila embryos. Cell 80, 747–756 (1995).

30 Asaoka-Taguchi, M., Yamada, M., Nakamura, A., Hanyu, K. & Kobayashi, S. Maternal Pumilio acts together with Nanos in germline development in Drosophila embryos. Nat Cell Biol 1, 431–437, doi:10.1038/15666 (1999).

31 Forstemann, K. et al. Normal microRNA maturation and germ-line stem cell maintenance requires Loquacious, a double-stranded RNA-binding domain protein. PLoS Biol 3, e236, doi:10.1371/journal.pbio.0030236 (2005).

32 Jin, Z. & Xie, T. Dcr-1 maintains Drosophila ovarian stem cells. Curr Biol 17, 539–544, doi:10.1016/j.cub.2007.01.050 (2007).

33 Park, J. K., Liu, X., Strauss, T. J., McKearin, D. M. & Liu, Q. The miRNA pathway intrinsically controls self-renewal of Drosophila germline stem cells. Curr Biol 17, 533–538, doi:10.1016/j.cub.2007.01.060 (2007).

34 Yang, L. et al. Argonaute 1 regulates the fate of germline stem cells in Drosophila. Development 134, 4265–4272, doi:10.1242/dev.009159 (2007).

35 Cox, D. N., Chao, A. & Lin, H. piwi encodes a nucleoplasmic factor whose activity modulates the number and division rate of germline stem cells. Development 127, 503–514 (2000).

36 Vagin, V. V. et al. A distinct small RNA pathway silences selfish genetic elements in the germline. Science 313, 320–324, doi:10.1126/science.1129333 (2006).

37 Ma, X. et al. Piwi is required in multiple cell types to control germline stem cell lineage development in the Drosophila ovary. PLoS One 9, e90267, doi:10.1371/journal.pone.0090267 (2014).

38 Ma, X. et al. Aubergine Controls Germline Stem Cell Self-Renewal and Progeny Differentiation via Distinct Mechanisms. Dev Cell 41, 157–169 e155, doi:10.1016/j.devcel.2017.03.023 (2017).

39 Maines, J. Z., Park, J. K., Williams, M. & McKearin, D. M. Stonewalling Drosophila stem cell differentiation by epigenetic controls. Development 134, 1471–1479, doi:10.1242/dev.02810 (2007).

40 Buszczak, M., Paterno, S. & Spradling, A. C. Drosophila stem cells share a common requirement for the histone H2B ubiquitin protease scrawny. Science 323, 248–251, doi:10.1126/science.1165678 (2009).

41 Eliazer, S., Shalaby, N. A. & Buszczak, M. Loss of lysine-specific demethylase 1 nonautonomously causes stem cell tumors in the Drosophila ovary. Proc Natl Acad Sci U S A 108, 7064–7069, doi:10.1073/pnas.1015874108 (2011).

42 Wang, X. et al. Histone H3K9 trimethylase Eggless controls germline stem cell maintenance and differentiation. PLoS Genet 7, e1002426, doi:10.1371/journal.pgen.1002426 (2011).

43 Xi, R. & Xie, T. Stem cell self-renewal controlled by chromatin remodeling factors. Science 310, 1487–1489, doi:10.1126/science.1120140 (2005).

44 James, T. C. & Elgin, S. C. Identification of a nonhistone chromosomal protein associated with heterochromatin in Drosophila melanogaster and its gene. Mol Cell Biol 6, 3862–3872 (1986).

45 James, T. C. et al. Distribution patterns of HP1, a heterochromatin-associated nonhistone chromosomal protein of Drosophila. Eur J Cell Biol 50, 170–180 (1989).

46 Eissenberg, J. C. et al. Mutation in a heterochromatin-specific chromosomal protein is associated with suppression of position-effect variegation in Drosophila melanogaster. Proc Natl Acad Sci U S A 87, 9923–9927 (1990).

47 Fanti, L., Giovinazzo, G., Berloco, M. & Pimpinelli, S. The heterochromatin protein 1 prevents telomere fusions in Drosophila. Mol Cell 2, 527–538 (1998).

48 Perrini, B. et al. HP1 controls telomere capping, telomere elongation, and telomere silencing by two different mechanisms in Drosophila. Mol Cell 15, 467–476, doi:10.1016/j.molcel.2004.06.036 (2004).

49 Piacentini, L., Fanti, L., Berloco, M., Perrini, B. & Pimpinelli, S. Heterochromatin protein 1 (HP1) is associated with induced gene expression in Drosophila euchromatin. J Cell Biol 161, 707–714, doi:10.1083/jcb.200303012 (2003).

50 De Lucia, F., Ni, J. Q., Vaillant, C. & Sun, F. L. HP1 modulates the transcription of cell-cycle regulators in Drosophila melanogaster. Nucleic Acids Res 33, 2852–2858, doi:10.1093/nar/gki584 (2005).

51 Vakoc, C. R., Mandat, S. A., Olenchock, B. A. & Blobel, G. A. Histone H3 lysine 9 methylation and HP1gamma are associated with transcription elongation through mammalian chromatin. Mol Cell 19, 381–391, doi:10.1016/j.molcel.2005.06.011 (2005).

52 Lin, C. H. et al. Heterochromatin protein 1a stimulates histone H3 lysine 36 demethylation by the Drosophila KDM4A demethylase. Mol Cell 32, 696–706, doi:10.1016/j.molcel.2008.11.008 (2008).

53 Piacentini, L. et al. Heterochromatin protein 1 (HP1a) positively regulates euchromatic gene expression through RNA transcript association and interaction with hnRNPs in Drosophila. PLoS Genet 5, e1000670, doi:10.1371/journal.pgen.1000670 (2009).

54 Kwon, S. H. et al. Heterochromatin protein 1 (HP1) connects the FACT histone chaperone complex to the phosphorylated CTD of RNA polymerase II. Genes Dev 24, 2133–2145, doi:10.1101/gad.1959110 (2010).

55 Xing, Y. & Li, W. X. Heterochromatin components in germline stem cell maintenance. Sci Rep 5, 17463, doi:10.1038/srep17463 (2015).

56 Zeng, A. et al. Heterochromatin protein 1 promotes self-renewal and triggers regenerative proliferation in adult stem cells. J Cell Biol 201, 409–425, doi:10.1083/jcb.201207172 (2013).

57 Abe, K. et al. Loss of heterochromatin protein 1 gamma reduces the number of primordial germ cells via impaired cell cycle progression in mice. Biol Reprod 85, 1013–1024, doi:10.1095/biolreprod.111.091512 (2011).

58 Brown, J. P. et al. HP1gamma function is required for male germ cell survival and spermatogenesis. Epigenetics Chromatin 3, 9, doi:10.1186/1756-8935-3-9 (2010).

59 Yan, D. et al. A regulatory network of Drosophila germline stem cell self-renewal. Dev Cell. 28, 459–473, doi: 10.1016/j.devcel.2014.01.020 (2014).

60 Rorth, P. Gal4 in the Drosophila female germline. Mech Dev 78, 113–118 (1998).

61 Dietzl, G. et al. A genome-wide transgenic RNAi library for conditional gene inactivation in Drosophila. Nature 448, 151–6, doi:10.1038/nature05954 (2007).

62 Petrella, L. N., Smith-Leiker, T. & Cooley, L. The Ovhts polyprotein is cleaved to produce fusome and ring canal proteins required for Drosophila oogenesis. Development 134, 703–712, doi:10.1242/dev.02766 (2007).

63 Lasko, P. F. & Ashburner, M. The product of the Drosophila gene vasa is very similar to eukaryotic initiation factor-4A. Nature 335, 611–617, doi:10.1038/335611a0 (1988).

64 Raz, E. The function and regulation of vasa-like genes in germ-cell development. Genome Biol 1, REVIEWS1017, doi:10.1186/gb-2000-1-3-reviews1017 (2000).

65 Hudson, A.M. & Cooley L. Methods for studying oogenesis. Methods 68, 207–217, doi: 10.1016/j.ymeth.2014.01.005 (2014).

66 Zhu, C.H. & Xie, T. Clonal expansion of ovarian germline stem cells during niche formation in Drosophila. Development 130, 2579–2588, doi: 10.1242/dev.00499 (2003).

67 Deng, W. & Lin, H. Spectrosomes and fusomes anchor mitotic spindles during asymmetric germ cell divisions and facilitate the formation of a polarized microtubule array for oocyte specification in Drosophila. Dev Biol 189, 79–94, doi:10.1006/dbio.1997.8669 (1997).

68 McKearin, D. & Ohlstein, B. A role for the Drosophila bag-of-marbles protein in the differentiation of cystoblasts from germline stem cells. Development 121, 2937–2947 (1995).

69 Ohlstein, B. & McKearin, D. Ectopic expression of the Drosophila Bam protein eliminates oogenic germline stem cells. Development 124, 3651–3662 (1997).

70 Cortes, A. et al. DDP1, a single-stranded nucleic acid-binding protein of Drosophila, associates with pericentric heterochromatin and is functionally homologous to the yeast Scp160p, which is involved in the control of cell ploidy. EMBO J 18, 3820–3833, doi:10.1093/emboj/18.13.3820 (1999).

71 Haynes, S. R., Johnson, D., Raychaudhuri, G. & Beyer, A. L. The Drosophila Hrb87F gene encodes a new member of the A and B hnRNP protein group. Nucleic Acids Res 19, 25–31 (1991).

72 Amero, SA. Elgin, SCR. Beyer, AL. A unique ribonucleoprotein complex assembles preferentially on ecdysone-responsive sites in Drosophila melanogaster. Genes Dev 5, 188–200, doi: 10.1128/MCB.13.9.5323 (1991).

73 Brimacombe, R., Stiege, W., Kyriatsoulis, A. & Maly, P. Intra-RNA and RNA-protein cross-linking techniques in Escherichia coli ribosomes. Methods Enzymol 164, 287–309 (1988).

74 Ule, J. et al. CLIP identifies Nova-regulated RNA networks in the brain. Science 302, 1212–1215, doi:10.1126/science.1090095 (2003).

75 Pek, J. W., Osman, I., Tay, M. L. & Zheng, R. T. Stable intronic sequence RNAs have possible regulatory roles in Drosophila melanogaster. J Cell Biol 211, 243–251, doi:10.1083/jcb.201507065 (2015).

76 Verrotti, A. C. & Wharton, R. P. Nanos interacts with cup in the female germline of Drosophila. Development 127, 5225–5232 (2000).

77 Epstein, A. M., Bauer, C. R., Ho, A., Bosco, G. & Zarnescu, D. C. Drosophila Fragile X protein controls cellular proliferation by regulating cbl levels in the ovary. Dev Biol 330, 83–92, doi:10.1016/j.ydbio.2009.03.011 (2009).

78 Xie, T. Control of germline stem cell self-renewal and differentiation in the Drosophila ovary: concerted actions of niche signals and intrinsic factors. Wiley Interdiscip Rev Dev Biol 2, 261–273, doi:10.1002/wdev.60 (2013).

79 Huertas, D., Cortes, A., Casanova, J. & Azorin, F. Drosophila DDP1, a multi-KH-domain protein, contributes to centromeric silencing and chromosome segregation. Curr Biol 14, 1611–1620, doi:10.1016/j.cub.2004.09.024 (2004).

80 Singh, A. K. & Lakhotia, S. C. The hnRNP A1 homolog Hrp36 is essential for normal development, female fecundity, omega speckle formation and stress tolerance in Drosophila melanogaster. J Biosci 37, 659–678 (2012).

81 Yan, D. & Perrimon, N. spenito is required for sex determination in Drosophila melanogaster. Proc Natl Acad Sci U S A 112, 11606–11611, doi:10.1073/pnas.1515891112 (2015).

82 Grieder, N. C., de Cuevas, M. & Spradling, A. C. The fusome organizes the microtubule network during oocyte differentiation in Drosophila. Development 127, 4253–4264 (2000).

83 Pisano, C. Bonaccorsi, S. Gatti, M. The kl-3 loop of the Y chromosome of Drosophila binds a tektin-like protein. Genetics 133, 569–579 (1993).

84 Moore, M. J. et al. Mapping Argonaute and conventional RNA-binding protein interactions with RNA at single-nucleotide resolution using HITS-CLIP and CIMS analysis. Nat Protoc 9, 263–293, doi:10.1038/nprot.2014.012 (2014).

85 Menet, J. S., Abruzzi, K. C., Desrochers, J., Rodriguez, J. & Rosbash, M. Dynamic PER repression mechanisms in the Drosophila circadian clock: from on-DNA to off-DNA. Genes Dev 24, 358–367, doi:10.1101/gad.1883910 (2010).

86 Livak, K. J. & Schmittgen, T. D. Analysis of relative gene expression data using real-time quantitative PCR and the 2(-Delta Delta C(T)) Method. Methods 25, 402–408, doi:10.1006/meth.2001.1262 (2001).

87 Jao, C. Y. & Salic, A. Exploring RNA transcription and turnover in vivo by using click chemistry. Proc Natl Acad Sci U S A 105, 15779–15784, doi:10.1073/pnas.0808480105 (2008).

