## Supplementary material for "Heterochromatin protein 1 (HP1) is intrinsically required for post-transcriptional regulation of *Drosophila* Germline Stem Cell (GSC) maintenance"

### Supplementary Methods

#### *Drosophila* Strains

The Ore-R stock and balancer stocks, used to balance inserts on the X, second, and third chromosomes, respectively, have been kept in our laboratory for many years.

The Gal4 lines used in this study were:

P{GAL4-nos.NGT}40 (Bloomington Drosophila Stock Center, BL #25751), P{matα4-GAL-VP16} (a gift from V. Palumbo) and P{bam promoter-Gal4:VP16}[17]. Flies carrying the UAS-mCD8-GFP reporter (BL#5130) were crossed with Gal4 drivers to check their expression pattern. *In vivo* RNAi experiments were performed at 25 °C.

HP1 RNAi lines, $w^{1118}$; P{GD 12524}v31994 and P{GD 12524}v31995 were obtained from the Vienna Drosophila Resource Center.

Flies carrying P{hs- bam.O}11d transgene (BL #24637) were used to generate *bam* ectopic expression by heat shock induction.

#### Immunofluorescent staining of larval, pupal and adult whole-mount ovaries

Pupal and adult ovaries were dissected in 1× PBS solution and fixed in 6% methanol-free formaldehyde (diluted in 1× PBS) for 10 min at room temperature. Larval ovaries were dissected in 1× PBS, freezing in liquid nitrogen and Methanol-Acetone fixed. The preparations were further permeabilized in 1× PBS containing 0.1% Triton (PBT), and then blocked for 20 min in blocking solution (1× PBS plus 5% not-fat dry milk).

The slides were incubated with primary antibodies overnight at 4 °C in a humid chamber. The following primary antibodies were used: chicken anti-GFP (Novus Biologicals), monoclonal mouse anti-HP1 9A9 (1:50, kindly provided by B. Wakimoto), rabbit anti-Vasa (1:50, Santa Cruz), mouse anti-α-Spectrin (1:50, DSHB), mouse anti-Bam (1:10, DSHB), rabbit anti-cleaved Caspase-3 (Asp175) (1:50, Cell Signaling Technology), rabbit anti-PH3 (kindly provided by P. Dimitri). After washing, the slides were incubated with secondary antibodies for 1 h at room temperature in a humid chamber.

Fluorescent labeled secondary antibodies raised in goat were used: FITC- coniugated anti chicken (1:50, Jackson ImmunoResearch), Alexa Fluor 568 coniugated anti mouse (1:300, Invitrogen), Cy2-linked anti mouse (1:50, Amersham), Alexa Fluor 555 coniugated anti rabbit (1:300, Invitrogen), Alexa Fluor 488 coniugated anti rabbit (1:300, Invitrogen).

Finally, the slides were washed three times in 1× PBS, stained with DAPI (4,6-diamidino-2-phenilindole, 0.01 mg/ml) or TOTO-3 Iodide (1μM) to visualize DNA and mounted in antifading medium (23,3 mg/ml of DABCO (Sigma Aldrich) in 90% glycerol/10% 1× PBS). All images were acquired on an Eclipse epifluorescence microscope (model E1000, Nikon) equipped with a CCD camera (Coolsnap) or Zeiss LSM 780 (Zeiss, Berlin, Germany) confocal microscope. Images were analyzed and further processed using Adobe Photoshop CS6.

#### Supplementary Figures and Figure legends

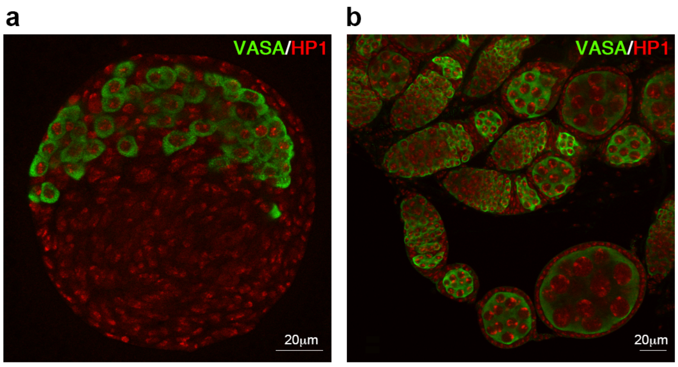


**Figure S1.** HP1 immunopattern on larval and pupal wild-type ovaries.

Confocal images of wild type larval (a) and pupal **(b)** ovaries double stained for anti-Vasa antibody (green) to label PGCs and germ cells and anti-HP1 antibody (red).

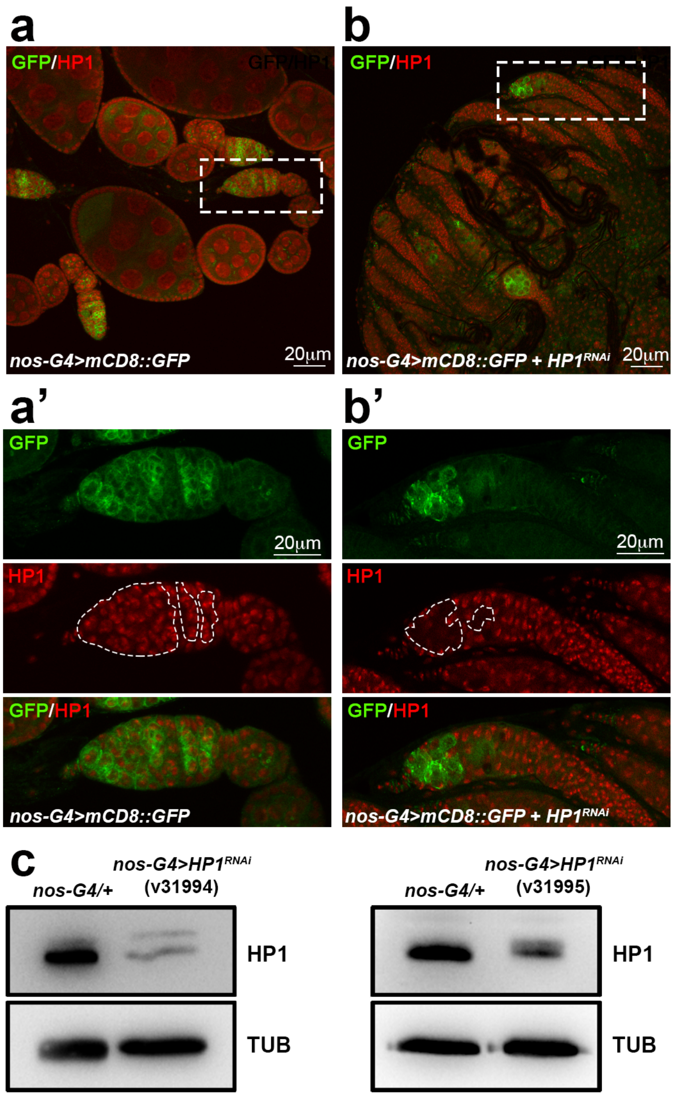


**Figure S2.** *In vivo* HP1 silencing by nos-Gal4 driver in adult ovaries. **(a)** Adult ovaries expressing mCD8-GFP driven by nos-Gal4 co-stained for HP1 (red) and GFP (green). **(a')** Magnified view of the dashed box in **(a)**. **(b)** *nos-Gal4>mCD8::GFP; HP1*$^{RNAi}$ ovaries stained for HP1 (red) and GFP (green). **(b')** Magnified view of the dashed box in **(b)**. HP1 staining is lost in germ cells marked by mCD8-GFP expressed by nos-Gal4 (outlined areas). **(c)** Western blot analysis showing HP1 protein levels in protein extracts from control ovaries (*nos-G4/+*) and HP1 depleted ovaries (*nos-G4>HP1*$^{RNAi}$) from two different HP1$^{RNAi}$ lines. α-Tubulin was used as a loading control.

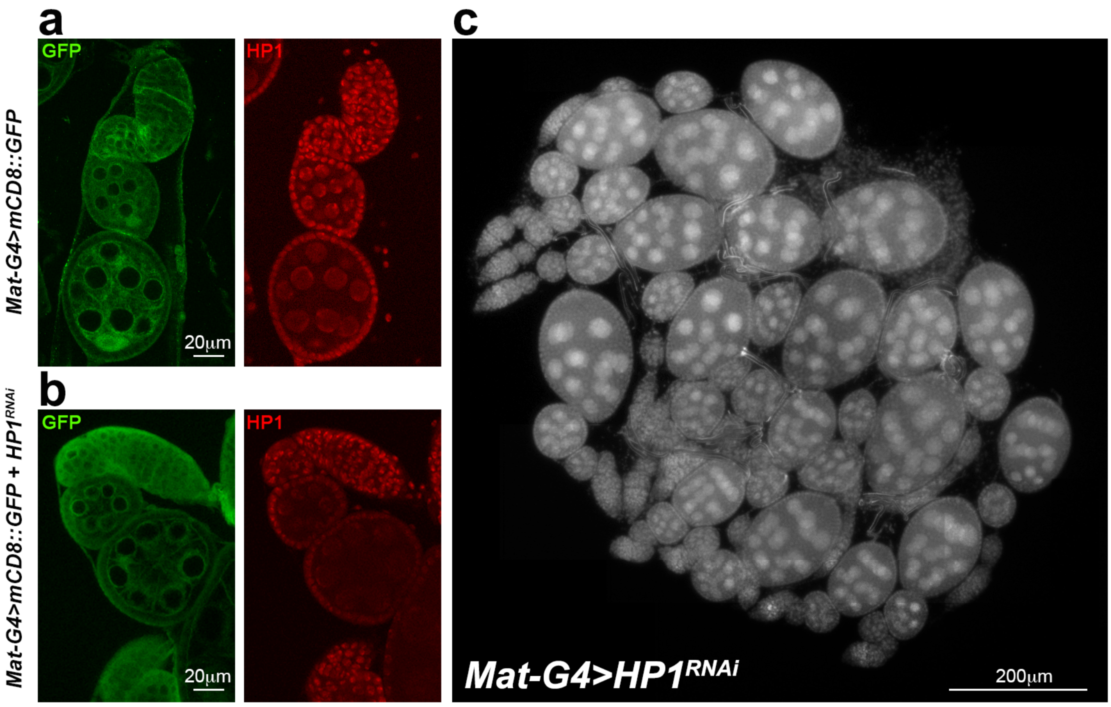


**Figure S3.** *In vivo* HP1 silencing by Mat-Gal4 driver. **(a)** Control (*Mat-Gal4>mCD8::GFP*) and **(b)** HP1 depleted ovaries (*Mat-Gal4>mCD8::GFP + HP1*$^{RNAi}$) stained for GFP (green) and HP1 (red). HP1 staining is lost in germ cells marked by mCD8-GFP expressed by Mat-Gal4. **(c)** DAPI staining of whole mount adult ovaries from *Mat-Gal4>HP1*$^{RNAi}$ females.

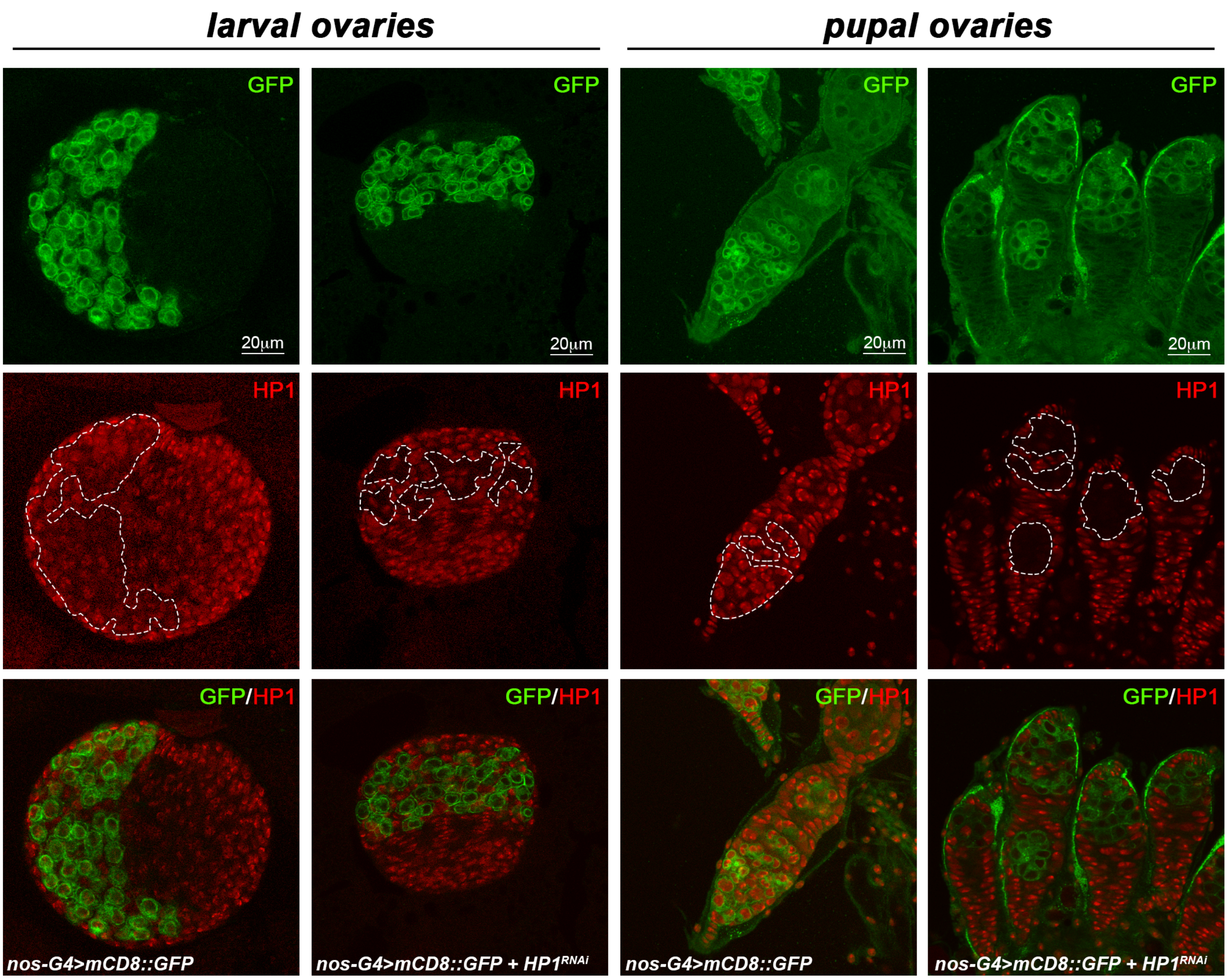


**Figure S4.** *In vivo* HP1 silencing by nos-Gal4 driver in larval and pupal ovaries.

Late third instar larval and pupal ovaries from *nos-Gal4> mCD8::GFP* and *nos-Gal4> mCD8::GFP +HP1$^{RNAi}$* females, co-stained for GFP (green) and HP1 (red). HP1 staining is completely lost in germ cells expressing mCD8 GFP by nos-Gal4 (outlined areas).

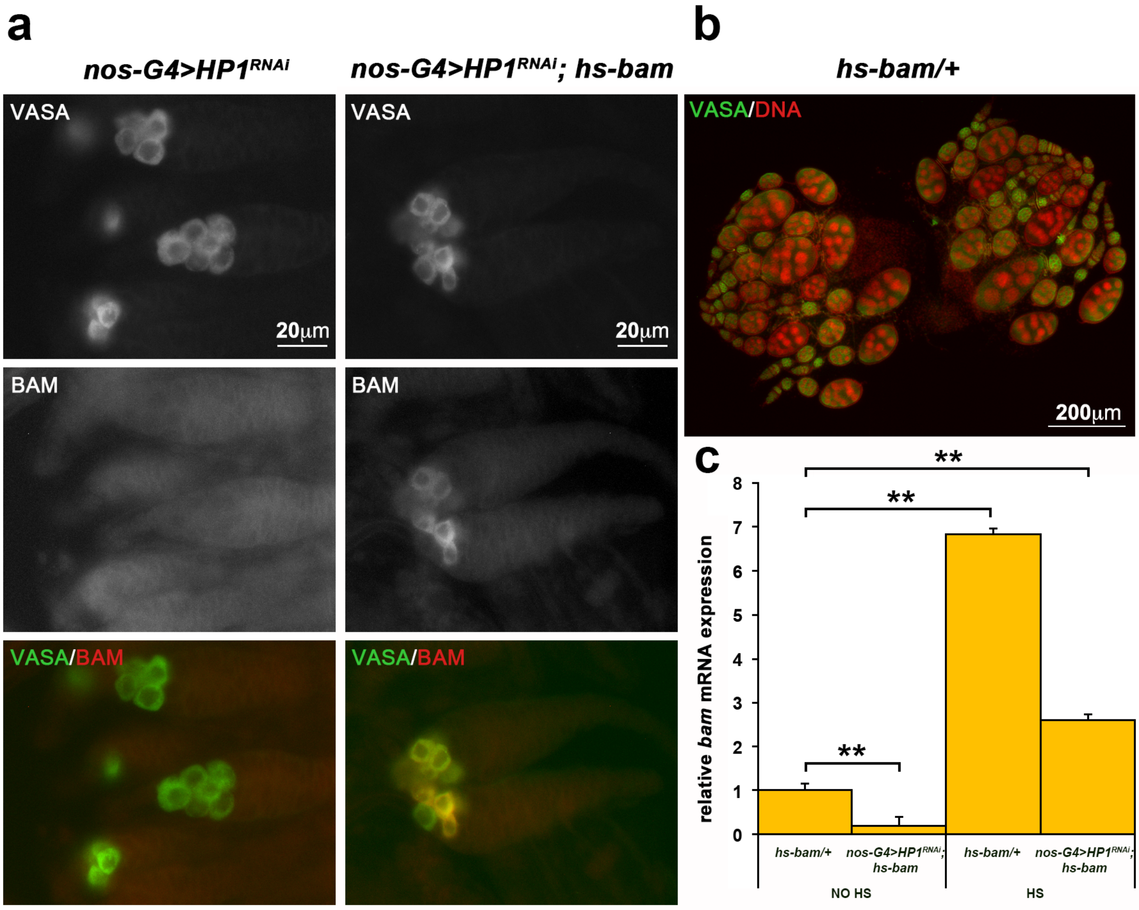


**Figure S5.** Ectopic expression of Bam in HP1 depleted ovaries. **(a)** Double immunostaining for Bam (green) and Vasa (red) on pupal ovaries from heat shocked (1 h at 37 °C followed by a 4 h recovery period before dissection) HP1 depleted females carrying (right panel) or not (left panel) the P[hs-bam] transgene. **(b)** Staining for Vasa (green) and DNA (red) on whole ovaries from heat shocked (1 h at 37 °C followed by a 24 h recovery period before dissection) control females carrying the P[hs-bam] transgene. The heat induced Bam expression itself does not alter the ovary morphology. **(c)** Relative *bam* mRNA expression level in control (*hs-bam/+*) and HP1 depleted ovaries (*nos-Gal4>HP1$^{RNAi}$; hs-bam*) from non heat shocked (NO HS) and heat shocked (HS, 1 h at 37 °C followed by a 2 h recovery period before dissection) females. The values shown are averages ±SEM of two biological replicates (***p* < 0.01).

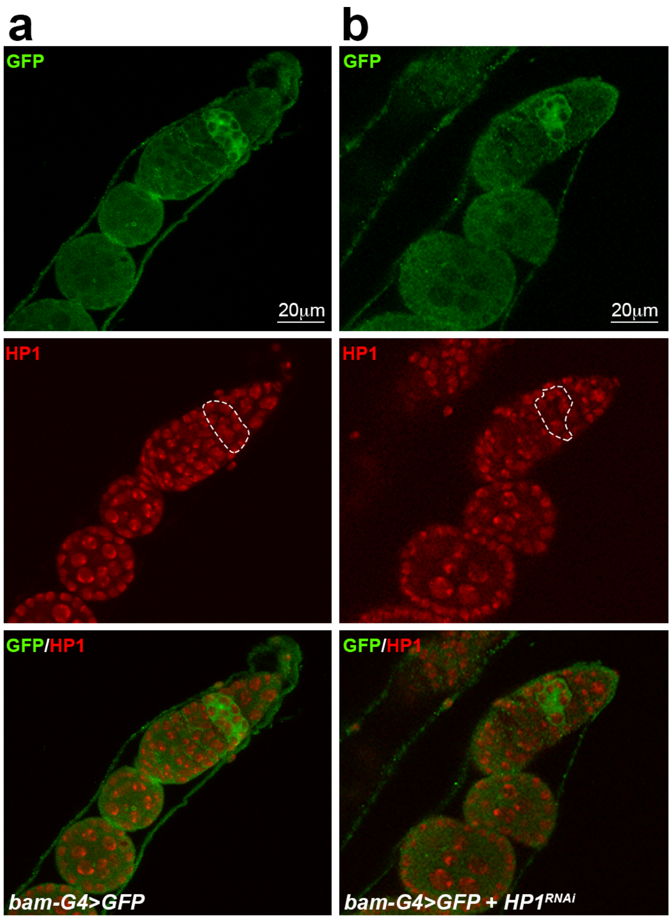


**Figure S6.** *In vivo* HP1 silencing by bam-Gal4 driver. **(a)** Control (*bam-Gal4> GFP*) and **(b)** HP1 depleted ovaries (*bam-Gal4> GFP + HP1*$^{RNAi}$) stained for GFP (green) and HP1 (red). The presence of HP1 immunosignals in GFP-positive cells (outlined areas) clearly indicates that bam-Gal4 expression is not sufficient to drive an efficient HP1 silencing.

**Supplementary Table S1: List of primers used in this study.**

| | Forward primer | Reverse primer |
|---|---|---|
| pum | GCCTGATGACCGATGTCTTTGG | CGATTTCCTGCTGCTGCTCC |
| mad | ACGAATCCTTCCAACAACTC | GAACACCCTTGCCTATATGAC |
| chico | CGCAGAGCAGCGATAAGCTA | GCGAACGCTCCAAGTTATCC |
| bam | TTGCTAATTGGTCTGCGCGATTGG | AACAGATCCTCCGCACTGATTCCA |
| tkv | GATTACCATTGCTGGTGCAAAGAAC | AAGAAGCCTCTTCGGTCGTAAAGAA |
| nos | CTGGCTCGATGCAGGATGTG | GTCTGCAGCTGGGCAGGATT |
| vas | CGGGACGACTTCTGGATTTC | GAGCAAGCTCCGCCTACAAT |
| medea | CGGCTATGTGGATCCCTCTG | CTAAGACAGCGCAGCCAGAC |
| effete | CAGTGGTCGCCAGCATTAAC | CACTCTCGTGCCAGCTCATT |
| fmr1 | CCTGGGAATAAGGCGGCACT | GCCTGCTGCTGTTGTACCTG |
| cup | CGACACCCAATTGCTACTGC | GGCTGCAAGAGTCTGCTGG |
| piwi | CAAGGCCGGATAATTGGAC | CCATCGCTCGGAGTGGTAAG |
| nos-prom | GGTGGTGTGCAGTGATTGTG | GGTAAAGCTACGCGCCAACT |
| bam-prom | TCAAATTTCTTTAAATGCGCCCGGG | GCCTTTTAAATCTTTCCTTATCGCCA |
